# Myelinated Inhibitory Axons in Human Neocortex

**DOI:** 10.1101/306480

**Authors:** Kristina D. Micheva, Edward F. Chang, Alissa L. Nana, William W. Seeley, Jonathan T. Ting, Charles Cobbs, Ed Lein, Stephen J Smith, Richard J. Weinberg, Daniel V. Madison

## Abstract

Numerous myelinated axons traverse the human neocortex. In a previous paper (Micheva et al., 2016) we showed that in mouse many of these axons belong to local inhibitory neurons, the parvalbumin-positive basket cells. Here, using samples of neurosurgically-excised cortex, we confirm the presence of myelinated inhibitory axons in all layers of human neocortex. As in mouse, these axons have distinctive features, including high neurofilament content, short nodes of Ranvier, and high content of myelin basic protein in their myelin sheath. We further show that, consistent with the known high-energy demands of parvalbumin interneurons, the inhibitory myelinated axons have more mitochondria, as well more 2’,3’-cyclic nucleotide 3’-phosphodiesterase (a protein enriched in the myelin cytoplasmic channels thought to provide access for trophic support from oligodendrocytes). The distinctive features of myelinated inhibitory axons in human cortical grey matter may have important implications for neurological disorders that involve pathologies of myelinated axons.

## Introduction

Myelinated axons, which account for approximately half the volume of the human brain (Filley & Fields, 2016; Wandell 2016), are a major factor enabling the dense, rapid, and efficient signal transmission that lies at the heart of human cognitive capacities. The insulating myelin sheath, created by a complex interaction between neuronal and glial cell, keeps axonal impulse propagation velocity high and energy consumption low, while allowing very small overall fiber diameters. Accordingly, when myelin is disturbed, brain function is profoundly affected, as observed in many degenerative neurological disorders, such as multiple sclerosis (Calabrese et al., 2015).

Myelinated axons convey information from a variety of neuronal types, and therefore the consequences of myelin abnormalities will vary depending on the neuronal type that is affected. However, very little is known about the diversity of central nervous system (CNS) myelinated axons, and the available evidence comes predominantly from studies in rodents (Gärtner et al., 2001; Jinno et al., 2007). Recently, we showed that a large fraction of myelin in the mouse neocortex ensheathes axons of inhibitory neurons, specifically of parvalbumin-positive basket cells (Micheva et al., 2016). These inhibitory myelinated axons differ significantly from the excitatory axons in structural organization (e.g., length of nodes of Ranvier and internodes), and in biochemical organization (e.g., cytoskeletal composition and protein content of myelin). Parvalbumin-positive myelinated axons have also been observed in human neocortex (Chung et al., 2013), and a recent study showed that the majority of cortical fast-spiking basket cells form myelinated axons (Stedehouder et al., 2017).

Do the inhibitory myelinated axons in human cortex share the distinctive features reported in mouse cortex? There are no available data regarding their relative abundance, laminar distribution, or possible differences with excitatory myelinated axons. In the present study we address these questions, using array tomography performed on surgically excised human neocortical tissue.

## Results and Discussion

The presence and distribution of myelinated axons was quantified in samples from human temporal neocortex. The tissue samples were obtained from surgical resections for epilepsy and prepared for array tomography (Micheva et al., 2010; Collman et al., 2015)(Table 1). Following resection, the tissue was immersion fixed with a mixture of paraformaldehyde and glutaraldehyde, and embedded in resin, similarly to Micheva et al., 2016. Serial ultrathin (70 nm) sections were cut from the tissue blocks, immunostained for the inhibitory neurotransmitter γ-aminobutyric acid (GABA), myelin basic protein (MBP), and other relevant markers, and imaged with a fluorescence microscope (Table 2).

**Table 1.** Human samples.

| Patient | Gender | Age | Clinical diagnosis | Region | Fixative |
| --- | --- | --- | --- | --- | --- |
| Q1010 | Male | 30 | Epilepsy | Anterior lateral temporal cortex | 2%PFA, 2%GA |
| Q1011 | Male | 44 | Epilepsy | Anterior medial temporal cortex | 4%PFA, 1%GA, 2.5% DMSO |
| Q1014 | Female | 48 | Epilepsy | Anterior lateral temporal cortex | 2%PFA, 4%GA, 2.5% DMSO |
| Q1015 | Female | 66 | Epilepsy | Anterior lateral temporal cortex | 2%PFA, 4%GA, 2.5% DMSO |
| Q1016 | Male | 49 | Epilepsy | Lateral temporal cortex | 2%PFA, 4%GA, 2.5% DMSO |
| Q1017 | Male | 46 | Epilepsy | Lateral temporal cortex | 2%PFA, 4%GA, 2.5% DMSO |
| Q1018 | Female | 20 | Epilepsy | Lateral temporal cortex | 2%PFA, 4%GA, 2.5% DMSO |
| Q1019 | Female | 51 | Epilepsy | Lateral temporal cortex | 2%PFA, 4%GA, 2.5% DMSO |
| 10/16/14 | Female | 30 | Nonspecific inflammation of the white matter | Frontal cortex | 4%PFA |

**Table 2.** Antibodies used in the study.

| Antigen | Host | Antibody Source | Dilution | RRID |
| --- | --- | --- | --- | --- |
| MBP | Chicken | AVES MBP | 1:200 | RRID:AB_2313550 |
| GABA | Guinea pig | Millipore AB175 | 1:5000 | RRID:AB_91011 |
| Parvalbumin | Rabbit | SWANT PV28 | 1:300 | RRID:AB_2315235 |
| Neurofilament heavy | Chicken | AVES NFH | 1:100 | RRID:AB_2313552 |
| α-Tubulin | Rabbit | Abcam ab18251 | 1:100 | RRID:AB_2210057 |
| Acetylated α-tubulin | Mouse | Sigma T6793 | 1:100 | RRID:AB_477585 |
| Glutamine Synthetase | Mouse | BD Biosciences 610517 | 1:25 | RRID:AB_397879 |
| PLP | Chicken | AVES PLP | 1:100 | RRID:AB_2313560 |
| CNPase | Chicken | AVES CNP | 1:100 | RRID:AB_2313538 |
| MDH2 | Rabbit | Origene TA308153 | 1:200 | RRID:AB_2722674 |
| TOMM20 | Rabbit | Abcam ab78547 | 1:100 | RRID:AB_2043078 |
| VDAC1 | Rabbit | ProteinTech 10866-1-AP | 1:50 | RRID:AB_2257153 |

MBP immunofluorescence clearly outlines myelinated axons, some of which contain the inhibitory neurotransmitter GABA (Figure 1). Myelinated GABA axons were particularly abundant in human cortical layers 3 and 4, and significantly sparser in layer 2. Myelinated GABA axons were rarely encountered in subcortical white matter. Overall, the proportion of myelinated axons that contain GABA was significantly lower in the human temporal cortex compared to mouse (somatosensory and visual cortex). For example, the highest percentage of GABA myelinated axons in human cortex was in layer 3a, 10 ± 2%, whereas GABA myelinated axons constitute 48 ± 3% of axons in mouse cortical layer 2/3 (Micheva et al., 2016).

**Figure 1.**
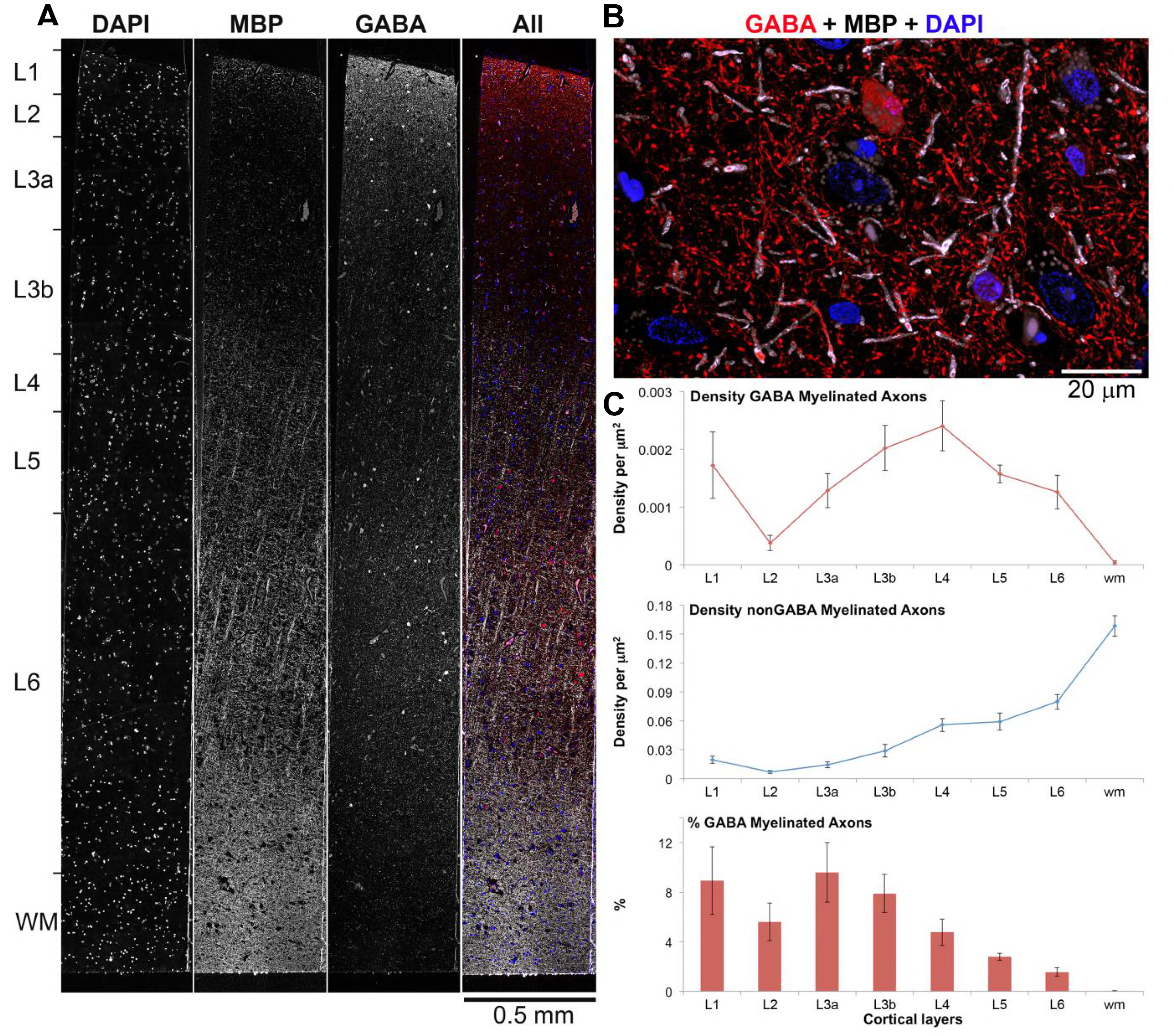
Distribution of inhibitory GABA myelinated axons in human temporal cortex. **A.** A 70 nm-thick section through human cortex immunostained with MBP (white) and GABA (red). Nuclei are labeled with DAPI (blue). **B.** Volume reconstruction of a subregion from layer 3a (35 serial sections, 70 nm each). **C.** Density and proportion of GABA myelinated axons in human temporal cortex. Means and standard errors from 8 human patients are shown.

The following source data are available for Figure 1:

**Source data 1**: Data values and statistics underlying Figure 1.

In our previous study of mouse cortex, we observed a number of differences between myelinated inhibitory and excitatory axons, which we now confirm in human cortex (Figure 2). Myelinated GABA axons in human cortex have significantly higher neurofilament content (average immunofluorescence of 819±55 a.u. vs 250±9 a.u. for non-GABA myelinated axons, p < 0.0001) and lower tubulin content compared to myelinated non-GABA axons (immunofluorescence of 233±19 vs 408±18 a.u., p < 0.0001; n= 141 GABA and 196 non-GABA myelinated axons from 3 different samples; Figure 2A-D). The nodes of Ranvier (Rasband & Peles, 2015), which appear as short gaps in the MBP and PLP staining along myelinated axons, are significantly shorter for myelinated GABA axons compared to unlabeled, presumably excitatory, axons (1.28±0.10 μm vs 1.79±0.07 μm, p < 0.0001; 39 GABA and 100 nonGABA nodes from 8 different samples; Figure 2F). Finally, GABA axons have more MBP in their myelin compared to neighboring non-GABA axons, immunofluorescence of 993±46 vs 854±30 a.u., p < 0.01; n= 141 GABA and 196 non-GABA myelinated axons from 3 different samples) even though their PLP content is essentially the same (1603±68 vs 1535±50 a.u., p = 0.41). Thus, the distinctive characteristics of myelinated inhibitory axons seen in mouse are also present in human cortex.

**Figure 2.**
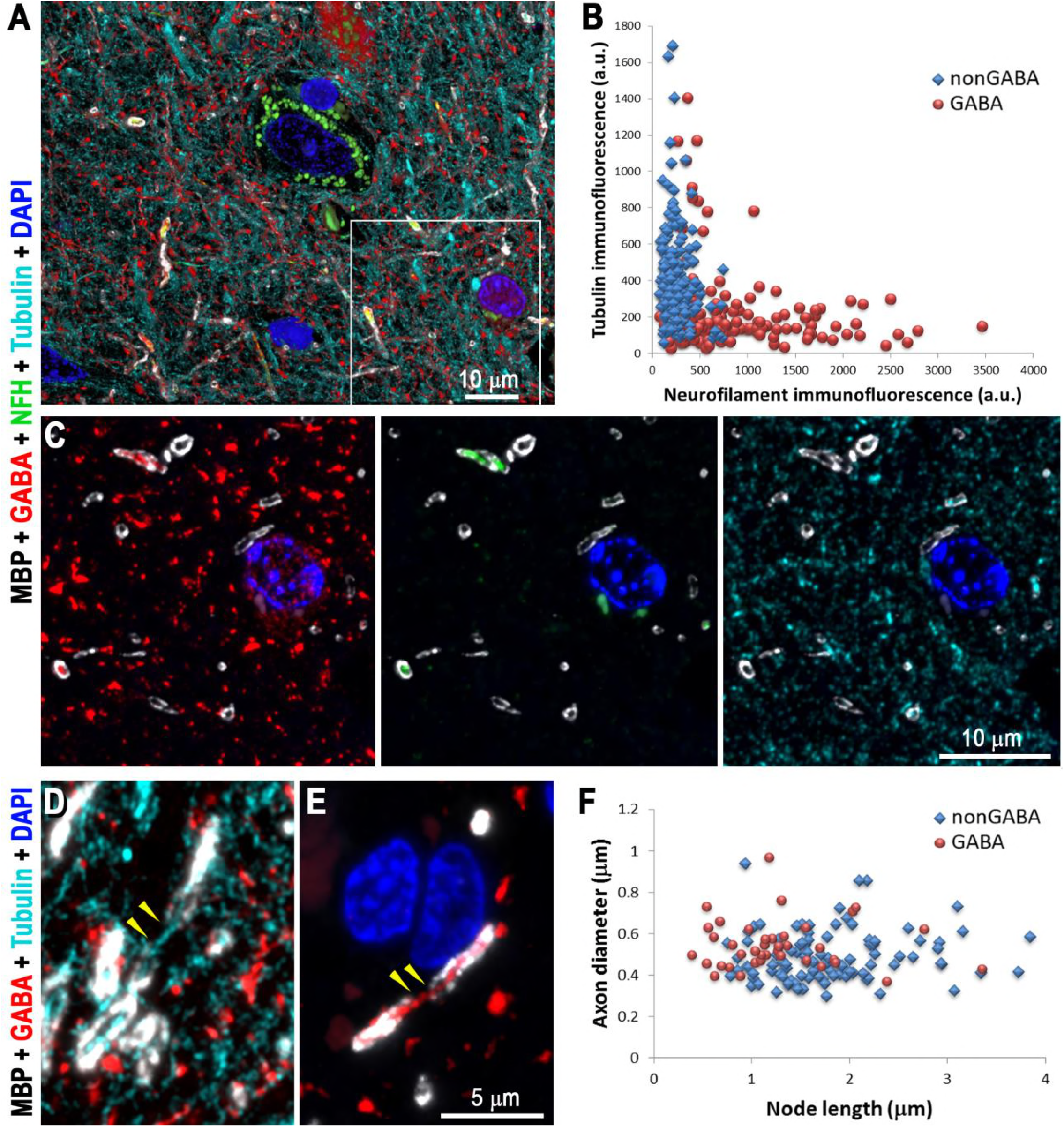
Myelinated GABA axons have distinct cytoskeletal composition and shorter nodes of Ranvier, compared to non-GABA myelinated axons. **A.** Volume reconstruction of 35 serial sections (70 nm each) from layer 3a of human cortex immunolabeled with GABA (red), MBP (white), neurofilament heavy chain (green) and α-tubulin (cyan). A single section from the boxed region is shown in C. **B.** Analysis of the cytoskeletal content of myelinated GABA vs. non-GABA axons from layers 3a, 3b and 4 from three human samples (196 non-GABA and 177 GABA myelinated axons). **C.** A single section from the boxed region in A, showing different combinations of immunostains. Note that myelinated GABA axons are brightly labeled with the neurofilament antibody, but have weak tubulin immunoreactivity. **D.** Maximum projection from 3 serial sections showing a node of Ranvier (yellow arrowheads) from a non-GABA axon. At the node, which is devoid of MBP immunofluorescence, the axon can be followed using the tubulin immunofluorescence (cyan). **E.** Node of Ranvier (yellow arrowheads) from a GABA axon. **F.** Comparison of the lengths of the nodes of Ranvier from GABA and non-GABA cortical axons (39 GABA and 100 nonGABA nodes, 8 samples).

The following source data are available for Figure 2:

**Source data 1**: Data values and statistics underlying Figure 2.

In mouse, the overwhelming majority of myelinated GABA axons come from parvalbumin-containing basket cells (Micheva et al., 2016). To determine the source of myelinated GABA axons in human cortex, the samples were also immunostained with an antibody against parvalbumin. More than half of the myelinated GABA axons were clearly immunoreactive for parvalbumin, with PV-immunofluorescent signal present on consecutive sections (Figure 3). On average, the PV immunofluorescence within GABA axons was significantly higher than within nonGABA axons (68±2 vs. 42 ± 1 a.u., p < 0.0001, n= 186 GABA and 172 non-GABA myelinated axons from 2 different samples). However, PV immunoreactivity in these samples was generally weak and displayed a low signal-to-noise ratio, making it difficult to determine the total number of PV-positive myelinated GABA axons. Low and variable levels of parvalbumin have been previously described in cortex of patients with epilepsy (Marco & DeFelipe, 1997).

**Figure 3.**
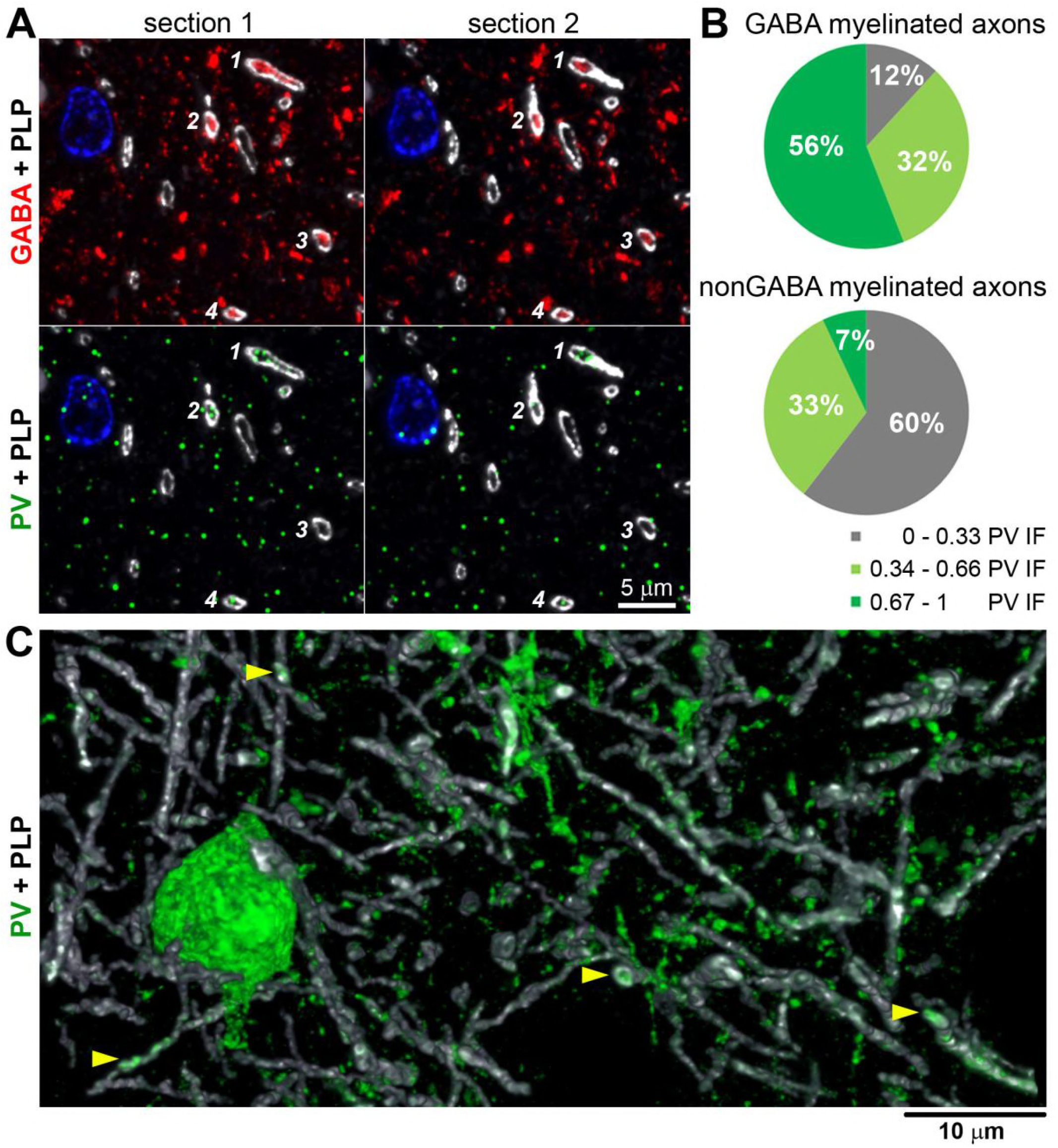
The majority of GABA myelinated axons are parvalbumin-immunopositive. **A.** Two consecutive sections from layer 3b of human cortex, immunolabeled for GABA (red), parvalbumin (green) and MBP (white). Cell nuclei are stained with DAPI (blue). Four myelinated GABA axonal profiles are marked with numbers; three of them (1, 2, and 4) are immunoreactive for parvalbumin. **B.** Pie charts showing the distribution of GABA and non-GABA myelinated profiles according to their parvalbumin immunoreactivity. Parvalbumin immunofluorescence was normalized for each sample (186 GABA and 172 non-GABA axonal profiles from two different samples). **C.** PV neuron and processes in layer 3b of human cortex from a biopsy - volume reconstruction from 49 serial sections. Yellow arrowheads point to PV-positive myelinated axons.

**Figure Supplement 1**. Cytoskeletal composition and node length of myelinated axons in epilepsy surgery vs. biopsy tissue.

The following source data are available for Figure 3:

**Source data 1**: Data values and statistics underlying Figure 3.

**Source data 2**: Data values and statistics underlying Figure Supplement 1.

To confirm our findings, we also used cortical tissue from a non-epileptic patient that had overlying cortical tissue removed during a biopsy to diagnose an inflammatory lesion. In this cortical specimen, we observe numerous myelinated axons immmunoreactive for parvalbumin. The proportion of the myelinated PV axons was very similar to the GABA axons in the tissue from epilepsy surgeries. For example, in layer 3a, 15% of the myelinated axonal profiles were PV immunopositive, and in layer 3b, 14% were PV immunopositive (693 myelinated axons from layer 3a and 2019 from layer 3b). This is within the range observed for GABA myelinated axons from epilepsy surgeries (3 - 21% in layer 3a, and 3-15% in layer 3b). The biopsy tissue, however, was fixed only with paraformaldehyde, which prevented simultaneous labeling with an anti-GABA antibody, since glutaraldehyde is required for satisfactory preservation of GABA (Somogyi et al, 1985; Schiffmann et al., 1988). Other characteristics of the PV-immunopositive myelinated axons from the non-epilepsy biopsy tissue, such as their cytoskeletal content and length of nodes of Ranvier (Figure 3 - Figure Supplement 1), were also very similar to the GABA axons in the epilepsy surgery tissue, suggesting that these represent comparable populations of axons. Thus, parvalbumin interneurons appear to be a major source of inhibitory myelinated axons in human neocortex.

Parvalbumin-positive basket cells are fast-spiking interneurons capable of generating long trains of action potentials at very high frequency (Kawaguchi et al., 1997; Kawaguchi & Kubota, 1997). The propagation of these action potentials along their axons requires a constant supply of ATP, and basket cells in rodent cortex are known for their high content of mitochondria (Gulyás et al., 2006). Accordingly, we observed significantly higher levels of immunofluorescence for three mitochondrial markers (MDH2, TOMM20, and VDAC1) in myelinated GABA compared to nonGABA myelinated axons, confirming that myelinated GABA axons in human cortex are enriched in mitochondria (Figure 4). How do these axons that are insulated by myelin receive the nutrients necessary for the mitochondrial production of ATP? Trophic support to myelinated axons in the CNS is likely provided through a system of cytoplasmic channels within myelin (reviewed in Simons & Nave, 2015), whose maintenance requires the myelin protein 2’,3’-cyclic nucleotide 3’-phosphodiesterase (CNP) (Snaidero et al., 2017). Our experiments reveal that GABA myelinated axons have a significantly higher CNP content compared to non-GABA axons (321±17 vs 271±7 a.u.; p = 0.001, 133 GABA and 489 non-GABA myelinated axons) suggesting the existence of more cytoplasmic channels within their myelin, consistent with a higher need for trophic support. Interestingly, this difference in CNP content between GABA and non-GABA axons is most pronounced among myelinated axons with a diameter larger than 0.6 μm (Figure 4K).

**Figure 4.**
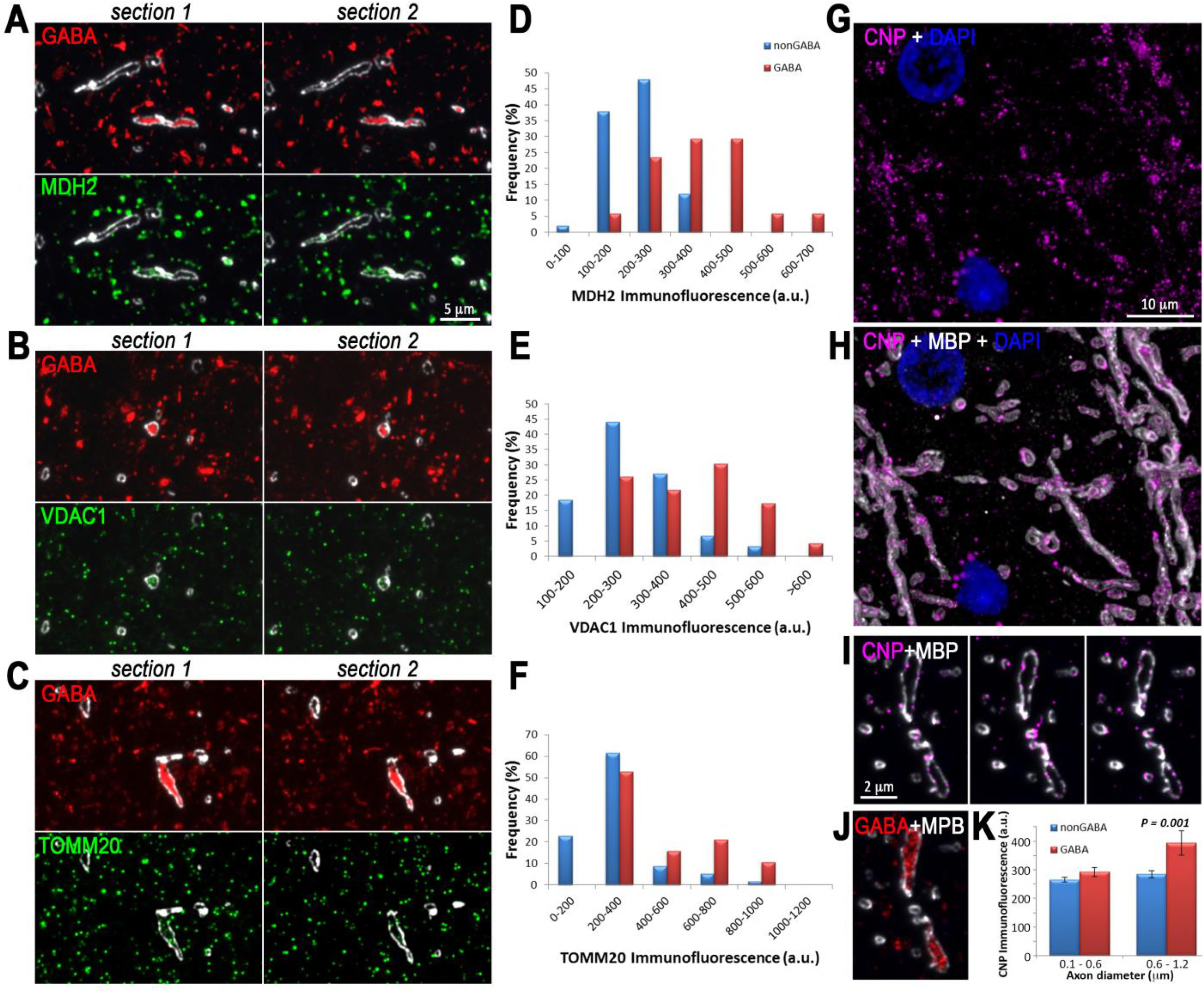
GABA myelinated axons have more mitochondria, and their myelin contains more CNP. **A-C.** Distribution of mitochondrial markers in myelinated axons, MDH2 (A), VDAC1 (B), and TOMM20 (C). In each case, two serial sections immunolabeled with GABA (red), MBP (white), and the mitochondrial marker (green) are shown. **D-F.** GABA myelinated axons have, on average, brighter immunofluorescence for MDH2, (D; data from 17 GABA and 60 non-GABA myelinated axons), VDAC1 (E; 23 GABA and 59 non-GABA) and T0MM20 (F; 19 GABA and 59 non-GABA). G,H, Volume reconstruction of 21 serial sections from human cortical layer 5, immunostained with CNP (magenta) and MBP. **I,** Three consecutive sections through a myelinated axons immunolabeled for CNP (magenta) and MBP (white). **J.** The first section from I, immunolabeled for GABA, shows that the myelinated axon in I is GABAergic. **K,** Thicker GABA axons have significantly more CNP in their myelin sheath compared to non-GABA axons of similar thickness (133 GABA and 489 non-GABA myelinated axons from 2 samples).

The following source data are available for Figure 1:

**Source data 1**: Data values and statistics underlying Figure 4.

Using samples of surgically-excised brain tissue, we show here that many of the characteristic features of myelinated axons in cortical grey matter previously described in the mouse (Micheva et al., 2016), are also present in human, despite 100 million years of evolutionary isolation. As in mouse, many human myelinated inhibitory axons originate from PV positive interneurons, and have distinctive features, including high neurofilament and low microtubule content, short nodes of Ranvier, and high content of MBP in their myelin sheath. The implications of these differences will require further study, but we speculate that they have direct functional significance. For example, the cytoskeleton is directly involved in axonal transport (Conde & Cáceres, 2009: Hirokawa et al., 2010); it also determines the mechanical properties of axons and therefore their susceptibility to mechanical injury (Ratzliff & Soltesz, 2000; Grevesse et al., 2015). The length of the nodes of Ranvier affects the speed and timing of action potential propagation (Arancibia-Cárcamo et al., 2017). MBP is a target in multiple sclerosis (Allegretta et al., 1990) and differences in its content may influence the fate of axons in brain pathologies. We further show that the inhibitory myelinated axons have more mitochondria, as well as more CNP, a protein enriched in the myelin cytoplasmic channels thought to provide access for trophic support from ensheathing oligodendrocytes. This is consistent with the high energy demands of fast-spiking PV interneurons and suggests that, in addition to influencing conduction velocity, the myelination of inhibitory axons is likely beneficial for managing their energy consumption by increasing the efficiency of action potential propagation (Hartline & Colman, 2007), and providing trophic support (Lee et al., 2012; Fünfschilling et al., 2012).

The characteristic features of myelinated inhibitory axons in human cortical grey matter are likely to have important implications for neurological disorders that involve pathologies of myelinated axons. The distinct molecular and structural organization of inhibitory and excitatory myelinated axons may underlie differences in their vulnerability in neurological disorders or to injuries. Furthermore, disturbances of myelin around inhibitory axons are expected to have very different functional implications compared to disturbances of myelin on excitatory axons and may require different strategies for prevention and treatment.

## Acknowledgements

This work was supported by NIH grants R01NS094499 to DVM, R01NS092474 to SJS, and R01NS039444 to RJW.

---

**Figure 3 - Figure Supplement 1.**
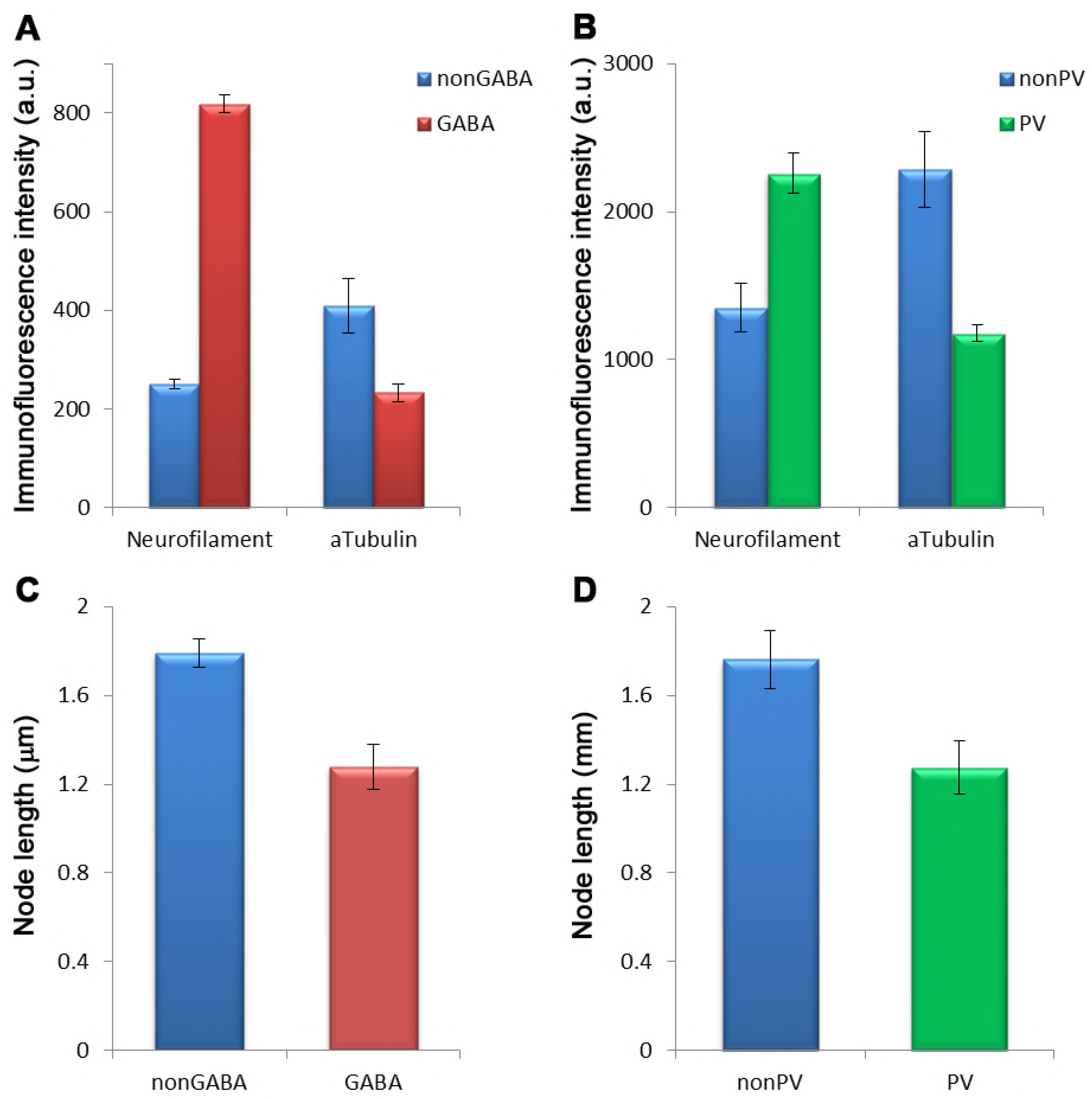
Cytoskeletal composition and node length of myelinated axons in epilepsy surgery vs. biopsy tissue. **A.** The immunofluorescence intensity (mean ± standard error) for neurofilament heavy chain and α-tubulin within myelin profiles is compared between nonGABA (blue) and GABA axons (red) from epilepsy surgery samples. The differences are statistically significant (p < 0.0001, t-test; 196 nonGABA and 177 GABA axons from layers 3a, 3b and 4 from three different samples). **B.** Comparison of the immunofluorescence intensity for neurofilament heavy chain and α-tubulin within myelin profiles of nonPV (blue) and PV axons (green) from the biopsy sample. The differences are statistically significant (p < 0.0001, t-test; 48 nonPV and 26 PV axons from layers 3a, 3b and 4 from one sample). **C.** Nodes of GABA axons are shorter than nonGABA axons in epilepsy surgery samples (p < 0.0001; t-test, 100 nonGABA and 39 GABA nodes from 8 different samples). **D.** Nodes of PV axons are shorter than nonPV axons in the biopsy sample (p = 0.01; t-test, 34 nonPV and 24 PV nodes).

